# *In vitro* Sex-specific Function-Structure Relationship in Neonatal Rat Cardiac Sheets

**DOI:** 10.1101/2025.07.10.664254

**Authors:** Mary Tran, Toby V. Nguyen, Suhani Khandelwal, Anna Grosberg

## Abstract

Even though *in vivo* rodent studies have been instrumental in investigating sex-specific differences in cardiac health, function, and pathology, they fall short in providing a fast and flexible platform for investigating sex differences of cardiac anisotropic monolayer in isolation. *In vitro* platforms offer an accessible and more controlled alternative to dissect and study the mechanisms by which male and female cardiac tissue sheets differ from one another. Here, we have shown on an *in vitro* heart-on-a-chip platform, primary neonatal rat ventricular myocytes can serve as a viable model showing sex chromosome-driven characteristics even in the absence of supplemented hormones. With controlled experimental conditions, the self-assembly of isolated cardiomyocytes resulted in morphological differences in the structure of the contractile apparatus. More importantly, the assembly of cardiac cells into confluent monolayers had a sex chromosome-driven divergence in both structure and the corresponding function. This work is the first characterization of the difference between sex-specific NRVMs in *in vitro* culture. Thus, this offers an avenue to investigate sex-based variations in cardiac function that are otherwise difficult to study.

## Introduction

Cardiovascular diseases, including heart failure, manifest differently in men and women, with distinct presentations and outcomes. As an example, women are more likely to develop heart failure with preserved ejection fraction, while men are more commonly affected by heart failure with reduced ejection fraction^1–3^, highlighting the divergence in cardiac physiology between the sexes. There is a wealth of knowledge on the pathophysiological differences between male and female hearts, encompassing both anatomical and functional aspects; and these dissimilarities likely originate from the dimorphism in the structure and function of the healthy organ. Indeed, men generally have larger and thicker-walled hearts^4–6^, and functionally, women’s hearts generate a smaller cardiac output at a faster rate and with larger ejection fraction^6–8^. These dissimilarities could potentially be influenced by any of the factors differentiating men and women, such as hormone variations, differences in cardiac structure and function, and distinct molecular pathways activated^1^. *In vivo* and *ex vivo* rodent studies have been instrumental in investigating sex-specific differences in heart failure, providing valuable insights into the physiological and pathological processes^9–12^. To highlight a few discoveries, these studies have shown that male rat cardiomyocytes are larger than females in all dimensions^13,14^, cardiac function is sexually dimorphic down to the myofibril with females exhibiting weaker and slower contraction kinetics^14–17^, and cardiac metabolism is a potential major contributor to sex dimorphism in heart failure pathogenesis. The divergence in metabolism occurs by differentially regulating the PKA pathway^18^, the mitochondrial function^19^, and metabolite utilization^20^. However, these *in vivo* and *ex vivo* models fall short in providing a fast and flexible platform for investigating specific aspects of sex differences in cardiac function and pathology. This gap can be partially addressed by utilizing *in vitro* engineered systems.

While there are many successful, functional *in vitro* cardiac tissues^21–37^, they often overlook sex as a biological variable or utilize induced pluripotent stem cells (iPSCs) derived cardiomyocytes, which are known to be immature and have high heterogeneity resulting from the differentiation process^38,39^. To engineer a more consistent cardiac tissue, some platforms use neonatal rodent ventricular myocytes, but many of these cardiac tissues have been generated using mixed-sex cells^22,23,27,28^. As a result, little is known about sex-specific structure and function of primary cardiomyocytes in engineered tissue, specifically of the intricate processes by which biological sex influences cellular organization and development, and the corresponding impact on overall cardiac function. This is a critical gap that needs to be addressed to make *in vitro* studies of sex differences more accessible, to develop more accurate models of heart disease, and to adapt existing *in vitro* platforms to study sex differences of the cardiac system.

Here, we utilized a previously developed heart-on-a-chip platform^40^ to produce sex-specific, 2D laminar cardiac sheets. We characterized primary neonatal rats’ sex chromosome-driven cell geometry and cytoskeletal self-assembly of their isolated cardiomyocytes in the absence of supplemented sex-specific hormones at the single cell level. Further, these confluent cardiomyocyte monolayers were analyzed both for structure and contractile functions. These findings will enable the augmentation of existing *in vitro* microphysiological systems, enabling them to support sex-specific ensembles, capturing their inherent differences, and providing a platform for testing targeted treatment strategies for cardiac injury as well as *in vitro* models for mechanistic studies.

## Results

Classically, primary *in vitro* cardiac tissues have been generated from mixed-sex ventricular myocytes with microcontact-printed fibronectin patterns of parallel stripes to promote anisotropic organization in the cardiac layer^33,40–43^. To explore the possibility of using these *in vitro* platforms to study sex differences, we characterized cardiac sheets engineered from neonatal rat ventricular myocytes (NRVMs) isolated from either female or male pups.

First, we sought to compare the structure of individual NRVMs when they are seeded sparsely onto a 20 *μ*m-wide line pattern. There were no qualitative differences between male and female NRVMs (Fig. 1(a)). Additionally, no significant difference in thickness was observed between the cells isolated from the two sexes (Fig. 1(b)). To quantify cytoskeletal architecture, cardiomyocytes were stained with *α*-actinin and actin, then analyzed using parameters previously developed to evaluate engieered cardiac monolayer tissue quality^44^. No statistically significant difference was found in the orientitation order parameter (OOP)(Fig. 1(c)), implying that the self-organization of cardiomyocyte contractile apparatus is independent of biological sex. However, there was a statistically significant reduction in both the continuous z-line length and z-line fraction (Fig. 1(d,e)) for the male cells compared to the female, suggesting a difference in the maturity level achieved by the self-assembly in isolation from other cell neighbors.

**Figure 1.**
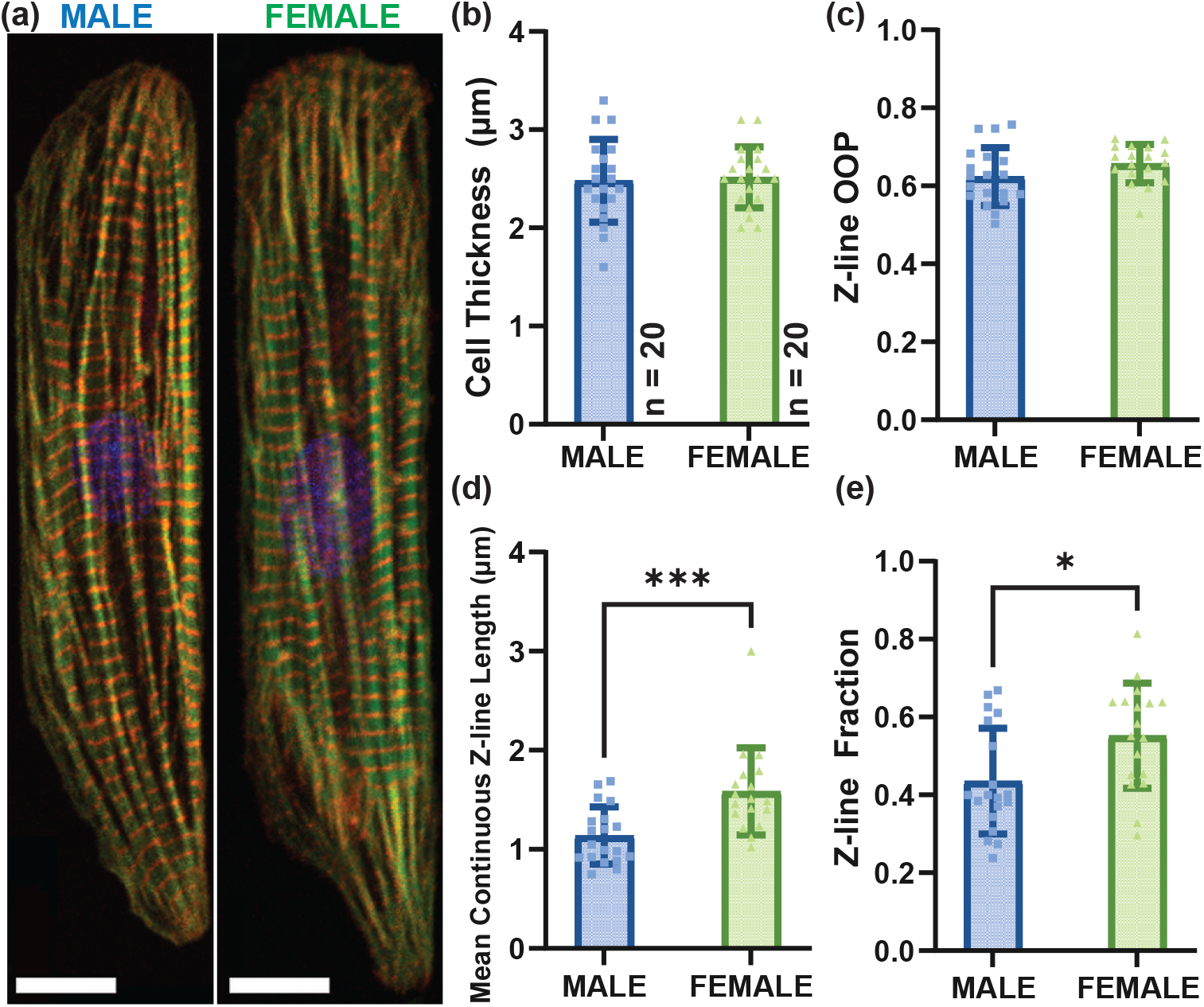
Single cell observations of sex-specific NRVMs. (a) Immunofluorescence of male and female individual cells stained for *α*-actinin (red), actin (green), and nuclei (blue). Scale bar = 10 *μ*m. (b) Thicknesses extracted from z-stacks of individual cells. (c,d,e) Sarcomeric structural metrics for isolated sex-specific cardiomyocytes. (c) Z-line organization order parameter (OOP) is a metric of degree of organized z-lines. (d) Mean continuous z-line is a metric for the quality of cardiomyocytes. (e) Z-line fraction is a metric of cell maturity. Each data point represents a singular cardiomyocyte, seeded from three separate litters for both female and male (Supplemental Table S1).

Next, the same metrics were used to characterized sex-specific self-assembly at a higher architectural level of confluent monolayers. Qualitatively, the myofibrils in female cultures were grouped in bundles, identifiable when they crossed (Fig. 2(a), many white triangles), while flat and sheet-like myofibrils were more characteristic of male NRVM cultures (Fig. 2(a), single white triangle). Corroborating with this observation, the self-assembly of these cardiomyocytes generated significantly thicker female NRVM sheets compared to their male counterparts (Fig. 2(b)). Z-line densities were then examined with respect to the total image area (Fig. 2(c)) and total cardiomyocytes area (Fig. 2(d)). There was a statistically significant difference between the two sexes for z-line density by image area (Fig. 2(c)), suggesting potential difference in the cell ensemble’s density given the same seeding density. However, neither sex had a significant advantage in cardiomyocyte z-line density (Fig. 2(d)). The sarcomeric architectures as described by previous metrics were not statistically different between male and female (Fig. 2(d)). Given all experimental conditions were controlled between the sexes, the observable differences suggested that the self-assembly of cardiac sheets was inherently sex chromosome-driven, but resulted in ensembles with similar z-line organization and maturation.

**Figure 2.**
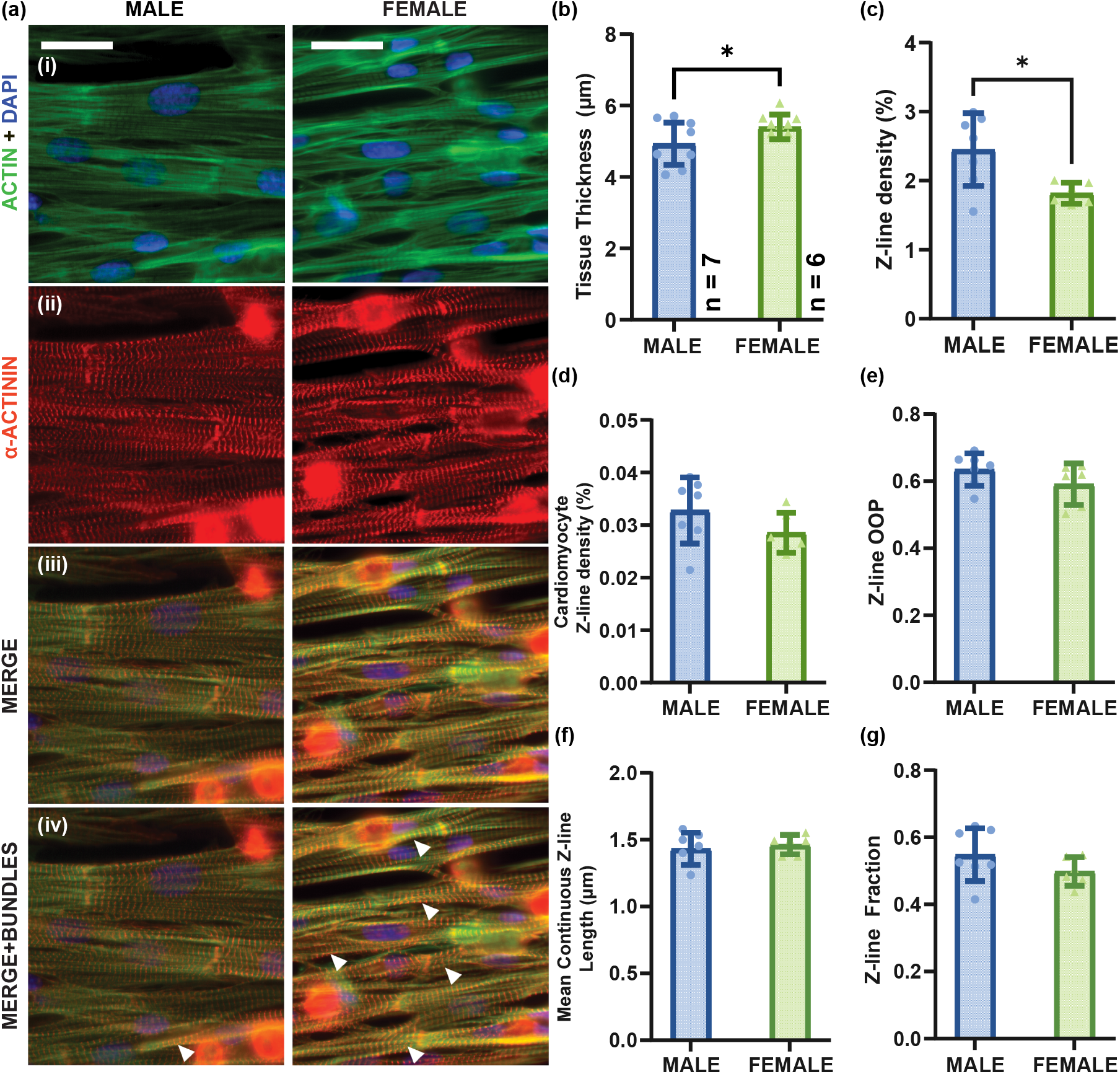
Sex-specific NRVM cardiac sarcomere structure. (a) Immunofluorescent staining of sex-specific NRVM sheets. Male NRVM cultures exhibited flat, sheet-like myofibril organization while female NRVMs form higher myofibril bundling (3D nature indicated by white arrows). Scale bar = 25 *μ*m. (b) Measured monolayer thicknesses. (c) Z-line density as represented by normalizing the total z-line pixels over total image area. (d) Cardiomyocyte z-line density as represented by normalizing the total z-line pixels over the total of actin pixels belonging to cardiomyocytes. (e,f,g) Sarcomeric structure metrics of female and male NRVM cultures^45^. (e) Z-line organization order parameter (OOP) is a metric of degree of organized z-lines. (f) Mean continuous z-line is a metric for the quality of cardiomyocyte structure. (g) Z-line fraction is a metric of maturity. Each data point represents a heart chip. The data in its entirety was analyzed from three litters of male and female pups each, with each litter producing 2 to 3 heart chips (Supplemental Table S1).

To explore functional differences between the sexes, NRVM sheets of varying densities (Table 1) were generated on the heart-on-a-chip platform as previously described^40^. The contraction of the cardiac films were recorded as stress traces over time, from which systolic, diastolic, and active stresses were extracted (Fig. 3(a)). As seeding density affected stress generation, systolic and active stresses were used to choose the optimal seeding density. Adjusted for the effects of the cells’ sex, there exists a trend of increasing systolic stress generation, statistically significant from low to mid seeding densities (generalized linear model, p = 0.027, 0.022, respectively, Supplementary Table S2). In active stress, the overall model did not find a significant effect of seeding density, but post hoc contrasts showed marginal differences within each group (Fig. 3(d)). Based on both findings, the mid density was chosen for both sexes for best performance.

**Table 1.** NRVM seeding density conditions used in heart-on-a-chip experiments. The sample size refers to the number of contracting films on the heart chip. For detail descriptions and animal allocation, see Supplemental Table S1.

| Density condition | # of cells seeded in 12-well plate | # of cells seeded per area | Sample sizes |  |
| --- | --- | --- | --- | --- |
| | (cells per well) | (cells per $\text{cm}^2$) | Male | Female |
| Low | $1.3 \times 10^6$ | $3.7 \times 10^5$ | 24 | 9 |
| Mid | $1.5 \times 10^6$ | $4.3 \times 10^5$ | 29 | 41 |
| High | $1.8 \times 10^6$ | $5.2 \times 10^5$ | 9 | 34 |

**Figure 3.**
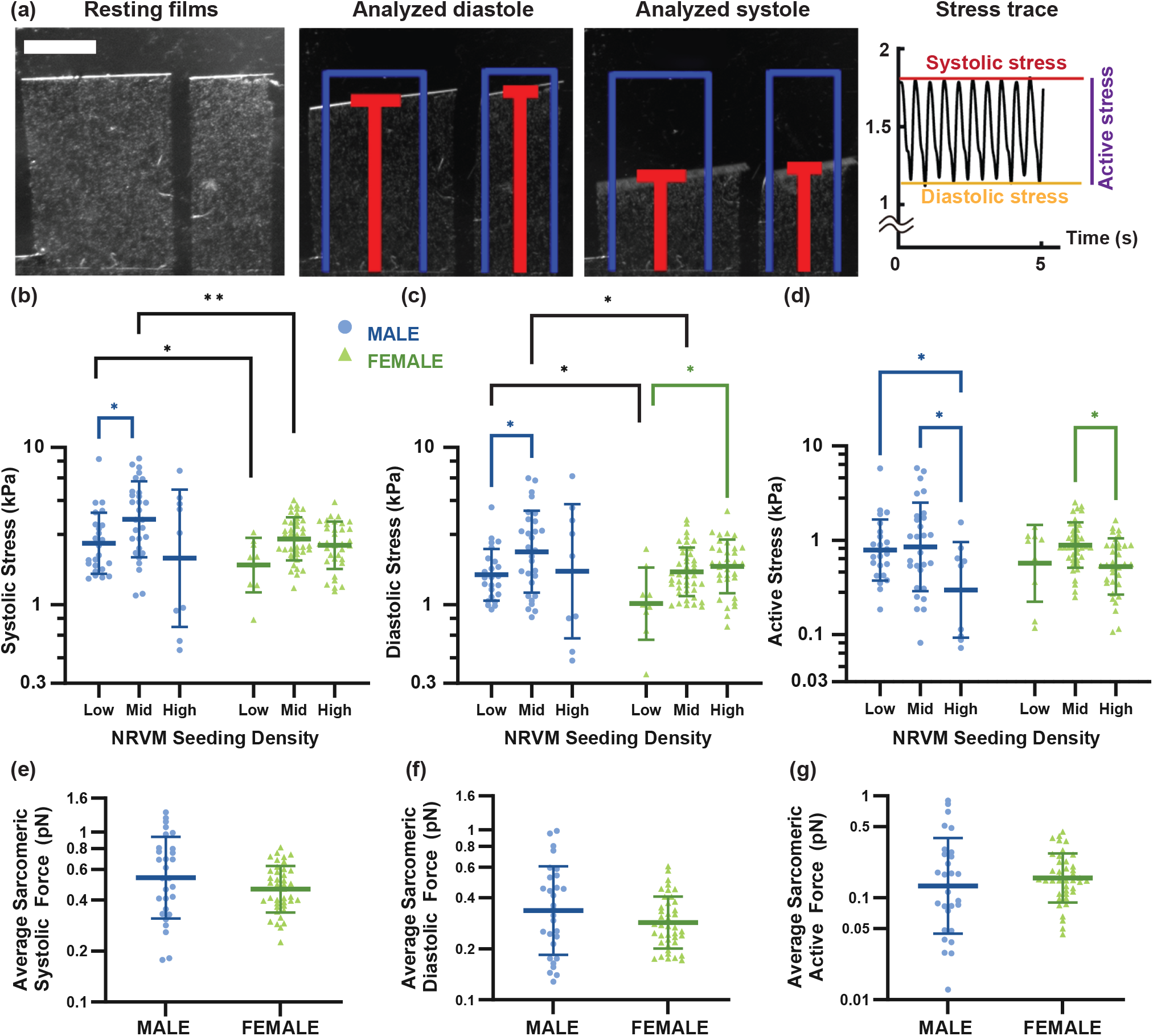
Contractility analyses. (a) Example heart-on-a-chip films at rest, at diastole, at systole, and the calculated stress trace over time. The width of the films do not influence stress calculation^32^. Scale bar = 1 mm. (b,c,d) Systolic, diastolic, and active stresses, respectively, of cardiac monolayers generated from sex-specific NRVMs at densities outlined in Table 1. (e,f,g) Average sarcomeric forces for systole, diastole, and active, respectively, computed by combining structural (cardiomyocyte area) and functional (stress generation) analyses at the mid seeding density (all stresses and forces are log-normally distributed. * p < 0.05, ** p < 0.01). Each data point represents a single film, with a range of 1 to 8 viable films per heart chip. The data was collected from 4 and 3 litters for male and female pups, respectively (Supplemental Table S1).

At the lower seeding densities (low and mid), male NRVM sheets were significantly stronger than females in generating both systolic and diastolic stress (Fig. 3(b,c)), showing a sex-dependent disparity in the ensembles’ abilities to contract and relax. However, no statistically significant sex-specific difference was observed in active stress generation (Fig. 3(d)), meaning the net stress produced was conserved between male and female cardiomyocyte ensembles. By combining the structural (Fig. 2(d)) and functional analyses, it was possible to estimate the average force for each sarcomere (Fig. 3(e,f,g)), which showed that at the micron scale, the two sexes were equivalent and matched previously estimated single sarcomere force^46,47^.

## Discussion

Our observations highlight that when male and female cardiomyocytes are exposed to identical experimental conditions— such as the same extracellular matrix pattern, culture media, and cell seeding density— their biological sex drives distinct morphological outcomes and functional profiles. At the single cell level, male and female NRVMs exhibited differences in their sarcomeric architecture (Fig. 1(d,e)), while maintaining comparable cellular height (Fig. 1(b)). The sex chromosome-driven, reciprocal effect between structure and function, then, emerged at a higher organization level. The self-assembly of confluent monolayers allowed the male cells to match with the female cells in terms of the z-line maturity metric (Fig. 1(c) vs. Fig. 2(c)) and z-line density within the cardiomyocytes area (Fig. 2(d)), which dovetails to the contractility results— where the active stress was preserved between both sexes (Fig. 3(b)). Finding the optimal contractility at the same seeding density is consistent with the consensus that the number of cardiac myocytes in the heart is the same in both sexes^5,48^. Yet, under these conditions, the male engineered NRVM sheets exhibited significantly elevated systolic and diastolic contractile forces compared to females at the same seeding density (Fig. 3(b,d, black)), which is in accordance with *ex vivo* observations^14–17^. Based on our estimate for average sarcomere force (Fig. 3(e-g)), the contribution of the individual sarcomeres to generate contraction is consistent between the two sexes and is independent of z-line structure and organization. Thus, the difference in functionality is due to the difference in sarcomeric structure observed between the two sexes after assembling into confluent layer, which resulted in the disparity in thickness and z-line density (Fig. 2(a-c)). The self-assembly process into these interconnected sheet itself, therefore, appears to be sexually dimorphic, driving male and female cardiomyocytes to adopt distinct architectures and impacting the emergent force production.

This sex difference demonstrated that *in vitro* neonatal rat ventricular myocytes can serve as a potential model for probing sex differences in cardiac research (Fig. 1,2,3). While there has been no direct measurement of human cardiac force, women exhibit lower end-diastolic and systolic pressure and volume^7,49^, as reflected in our neonatal rat model (Fig. 3(b-d)). These observations between rat and human cardiac characteristics suggest that primary rat neonatal cardiomyocytes provide a promising and accessible platform to explore sex-specific cardiac function, offering insights that may be applicable to human health. While adult cardiac physiology is shaped by both chromosomal and hormonal factors, the understanding of the relative contributions of each remains incomplete^18^. By isolating the effects of chromosomal sex in a simplified *in vitro* model, our work provides a foundational perspective that can inform and complement future studies incorporating hormonal influences or adult cardiomyocyte systems.

Indeed, even though NRVMs are developmentally immature at the time of isolation, engineered constructs similar to ours have been shown to have elongated morphology, aligned sarcomeres with appropriate lengths^50^(Supplemental Fig. S2), contractile function, inotropic response to changes in extracellular Ca^2+^ concentration, and gene expression profiles comparable to those of adult rat myocardium^51^, as well as slower spontaneous kinetics in females (Supplemental Fig. S3) as seen in isolated cardiomyocytes^15,16^, supporting the physiological relevance of this approach. NRVMs’ accessibility, cost-effectiveness, and rapid experimental turnaround make them particularly valuable for studies requiring high-throughput tissue generation and functional assessment, which means they remain a widely utilized model in cardiac research^52–57^. Nonetheless, NRVMs do not fully replicate the phenotypic complexity of adult human cardiomyocytes, and findings from this model should be interpreted within that limitation. Additionally, the native myocardium is a three-dimensional structure with complex fiber orientations—spiraling and tangentially aligned layers—as well as contributions from multiple cardiac cell types. In that sense, our current model represents a simplified version of cardiac tissue, consisting of a single anisotropic monolayer composed solely of ventricular myocytes. However, we emphasize that the platform’s modularity allows for added complexity. Recent work from our group has extended this system to incorporate macrophages and hypoxic injury, enabling investigation of immune–cardiac interactions^58^. These developments underscore the adaptability of the platform and its potential to model increasingly sophisticated aspects of cardiac physiology and disease, now with sex as a biological variable.

Many *in vitro* platforms have successfully used mixed-sex rat cardiomyocytes to generate functioning tissue^22,23,27–30^, which have been contributory to cardiovascular research. In this study, we demonstrated that sex-separated NRVMs can also serve as a valuable model for investigating sex dimorphism in cardiac biology. Beyond identifying structural and functional differences between male and female confluent monolayers, we established a link between these differences, providing insight into their cause-and-effect relationship in the absence of supplement hormones. This finding opens the door for future studies to explore the specific biological pathways that drive sex-based disparities in cardiac tissue, including those governed by sex hormones. Notably, we observed that these sex chromosome-driven differences became evident when cardiomyocytes self-assembled into organized monolayers, underscoring the importance of studying cardiac function at the tissue level rather than just isolated single cells. Furthermore, existing tissue engineering techniques can be readily adapted to support sex-specific rodent cardiac sheet, ensuring their applicability in modeling sex-specific cardiac function. This adaptability strengthens the utility of rodent cardiomyocytes as a robust platform for drug testing, disease modeling, and the development of targeted therapeutic strategies.

## Methods

### Substrate Preparation

The chips were fabricated as previously described and used^40^, with some modifications. Briefly, a large glass coverslip (76 mm x 83 mm; Brain Research Laboratories, Newton, MA) was cleaned via sonication for 10 minutes in 200-proof Ethanol and air dried. Protective film strips were placed parallel to the edge of the glass, approximately 1 cm apart. Next, poly(N-isopropylacrylamide), (PIPPAAm, Sigma-Aldrich, Burlington, MA) was dissolved in 99.4% 1-butanol at 10%wt (weight/volume). An excess of the PIPPAAm solution (1 mL total) was deposited and spin-coated onto the glass surfaces not covered by the protective films at 6000 RPM for 1 minute. The protective film was removed. Then, Sylgard 184 polydimethylsiloxane (PDMS; Ellsworth Adhesives, Germantown, WI) elastomer was mixed at a 10:1 base to curing agent ratio and partially cured at 37°C between 90 to 105 minutes prior to spin coating on the glass substrate, with a terminal velocity of 4000 RPM for 1 minute. Next, the glass substrate was left to cure overnight at 65°C. After curing, the PDMS was scored lightly to create 4 strips of thin films without engraving the glass underneath, and the glass substrate was then cut into individual chips with a laser cutter (Trotec Speedy 360; Trotec Laser GmbH., Plymouth, MA). Three chips from each large glass coverslip were saved for thickness measurements. PDMS film thicknesses were measured to be within a range of 12 to 15 microns using a profilometer (Dektak 6M, Veeco Instruments Inc., Plainview, NY).

Prior to seeding cells, fibronectin (FN; Sigma-Aldrich, St. Louis, MO) was microcontact-printed onto the surface of the PDMS substrate. The stamps were patterned to be parallel lines, 22 *μ*m in width with 3 *μ*m gaps in between, made from PDMS. The stamps were sterilized in 200-proof Ethanol, and let dry in a biohood under sterile conditions. The patterned surface of the dry stamps was covered in 200 uL of 50 *μ*g/ mL FN and incubated for 1 hour. The surface of the cover slips was sterilized and functionalized by exposing them for 8 min to UV ozone (Model No. 342, Jetlight Company, Inc., Irvine, CA). The stamps were dried with compressed nitrogen and used to transfer the FN pattern to the coverslips with 4 minutes of contact. A 12-well plate was used to house the chips, initially coated with 1% Pluronics F127 (BASF Group, Parsippany, NJ) in DI water for ten minutes and washed three times with Phosphate Buffered Saline (PBS; ThermoFisher, Waltham, MA, Cat#10010049). The chips were transferred to the Pluronics-coated plate, submerged in PBS, and stored at 4°C before use. All chips were used within 4 to 7 days after FN stamping. Chips used for structural assays were fabricated similarly^40^, without coating PIPPAAm nor scoring PDMS into films.

### Isolation and culture of neonatal rat ventricular myocytes (NRVMs)

All animal procedures were performed with the approval of the Institutional Animal Care and Use Committee of University of California, Irvine (IACUC Protocol #2022-054). The data was collected over seven and six separate isolation procedures for male and female experiments, respectively. Each isolation procedure used a litter of ten rat pups. All procedures and data analysis methods in this work adhered to the ARRIVE and other relevant guidelines and regulations.

The isolation of primary ventricular cardiomyocytes were performed as previously described^40^. Briefly, the provider Charles River Laboratories (Wilmington, MA) sexed two-day old neonatal Sprague-Dawley rats by the anogenital distance from multiple litters and provided groups of ten male-only or female-only pups with a random dam. Dams were euthanized with carbon dioxide asphyxiation followed by bilateral pneumothorax. The ventricles were extracted from only the pups, homogenized by washing in Hanks Balanced Salt Solution (HBSS; ThermoFisher, Waltham, MA, Cat#14170161), then incubated in 1 mg/mL trypsin solution (Sigma-Aldrich, St. Louis, MO) dissolved in HBSS for 12 hours at 4°C on a rocker. The trypsin was neutralized with M199 culture medium (Thermo Fisher, Waltham, MA, Cat#11150067) supplemented with 10% fetal bovine serum (FBS; ThermoFisher, Waltham, MA, Cat#26140079) at 37°C. The tissues were dissociated to single cells four times via digestion with 1 mg/mL collagenase type II solution (Worthington Biochemical Coporation, Lakewood, NJ) dissolved in HBSS at 37°C. Cells were washed in chilled HBSS, centrifuged at 1000 RPM for 6 minutes. Cells were then resuspended in warm M199 culture medium supplemented with 10% FBS, 10 nM HEPES, 3.5 g/L Glucose, 2 nM L-glutamine, 2 mg/L vitamin B-12, and 50 U/mL penicillin. The myocytes were purified by three repeats of differential plating at 37°C for one hour each. Each harvest, consisting of a litter of 10 pups, produced roughly 10 heart chips, which were split between structural and functional chips. Purified cells were then seeded on the heart chips at densities of 1.3 × 10^6^, 1.5 × 10^6^, or 1.8 × 10^6^ cells per well for confluent monolayers and at 1.5 × 10^5^ cells per well for single cells. Chips used in structure analyses were seeded at 1.5 × 10^6^ cells per well. The chips were cultured at 37°C and 5% CO_2_ over a period of four days prior to running the assays. During the initial 48 hours, the cells were kept in previously described media, supplemented with 30% FBS, and media was refreshed every 24 hours. After which the cells were incubated in 10% FBS supplemented media for the remaining 48 hours.

### Cardiac Structure Assay and Analysis

On day 5 of culturing the cardiomyocytes, the chips were rinsed with PBS prior to being fixed and permeabilized in a solution of 4% paraformaldehyde (PFA; VWR, Radnow, PA) and 0.05% Triton-X (Sigma-Aldrich, St. Louis, MO) in PBS for 10 minutes. The NRVM cultures were stained for actin (Alexa 488 Phalloidin, ThermoFisher, Waltham, MA), nuclei (4’,6-diamidino-2-phenylindole hydrochloride, DAPI; Invitrogen, Waltham, MA), sarcomeric *α*-actinin (clone EA-53, mouse monoclonal anti- *α*-actinin, Sigma-Aldrich,St. Louis, MO), and fibronectin (rabbit anti-fibronectin; Invitrogen, Waltham, MA) at a dilution of 1:200 for each antibody for 1 hour at room temperature. Secondary antibody solution was prepared with goat anti-mouse IgG (AlexaFluor594; ThermoFisher, Waltham, MA) and goat anti-rabbit IgG (Cy5; ThermoFisher, Waltham, MA) at 1:200 dilution, in which the chips were incubated for 1 hour at room temperature. Finally, the chips were mounted on glass slides with ProLong Glass Anti-fade Mountant (ThermoFisher, Waltham, MA).

### Image Acquisition

For cardiac monolayer structure images, imaging was performed on a digital CCD camera ORCA-R2 C10600-10B (Hamamatsu Photonics, Shizuoka Prefecture, Japan) mounted on an IX-83 inverted motorized microscope (Olympus America, Center Valley, PA). For thickness measurements, a laser scanning confocal microscope (Olympus Fluoview FV3000; Olympus America, Center Valley, PA) was used. All images were taken at 40X magnification. Ten randomly selected fields of view for each chip were obtained. Z-stacks were obtained at the top, middle, and bottom slices for NRVM sheets at ten random fields of view. Z-stacks for single cells were obtained at a resolution of 0.1 *μ*m and processed in Fiji to obtain thicknesses.

### Image Analyses

The cardiomyocytes and cardiac monolayer structures were quantified using a custom MATLAB script, as previously developed^45^. Binary skeletons of *α*-actinin signals were extracted from microscopy images and analyzed for z-line orientation order parameter (OOP), z-line fraction, and mean continuous z-line lengths as averages from the ten fields of view per chip.

To quantify cardiomyocyte area from imaging data, a decision tree classifier was implemented using a custom MATLAB script, as previously described^59^. A subset comprising approximately 1% of the dataset was manually labeled to identify cardiomyocyte regions, with a final distribution of 57% male and 43% female images. The classifier, optimized on this training set, achieved an accuracy of 98%. This classification reported the number of pixels of actin signals within the cardiomyocyte area for each sex.

### Contractility Assay

Contractility experiments were performed as previously described^40^ with some modifications. The chips were transferred from the incubator to a stereoscope in a 60 mm petri dish filled with warm, 37°C, media. To release the films, a razor blade was used to make two cuts in the middle of the chip, perpendicular to the PDMS scored lines, 0.5 to 1 mm apart. The resulting thin strip in between was peeled off with tweezers. Field stimulation electrodes were built from 1 mm diameter carbon rods (McMaster-Carr, Douglasville, GA) and attached to a PDMS mount in a parallel configuration where they were placed 1.5 cm apart. During the experiments, the PDMS mount was slotted into the edge of the 35 mm pertri dish containing the chip and warm, fresh media, with the electrodes pointing downwards and submerged into the solution. The chip was imaged between the electrodes. The chips were paced at 2 Hz, 20 V using an external field stimulator (Myopacer, IonOptix Corp., Milton, MA), which applied a square wave pulse. Video recordings of the films contracting captured at least 5 cycles for 1 Hz and 10 cycles for 2 Hz. All images and movies were collected using a Basler camera (A602f Basler Inc, Exton, PA) controlled by MATLAB, with all movies containing 100 frames per second. An image of a scale ruler was taken afterwards. Then the media was removed from the petri dish, allowing the films to lay flat on the glass surface. An image of the films was taken for length measurements, and another image of the scale ruler was taken to account for changes in focus. Contractility videos were analyzed for stress produced by the cardiac films using custom Fiji and MATLAB scripts as described^40^. Each chip produced a range of 1 to 8 viable films for analyses, with viable meaning the films were contracting spontaneously and were responding to the electrical stimulation.

The average sarcomeric force was determined by combining structural data of Normalized Cardiomyocyte Area and contractility data 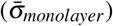:

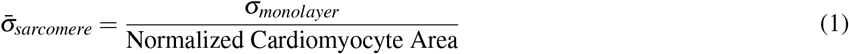

where,

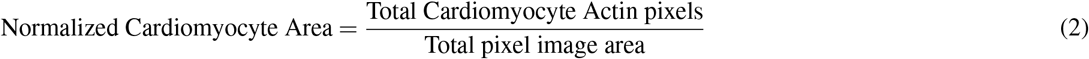

Then, using previously published^47^ estimate of sarcomere area, *A*_*sarcomere*_ = *π*(60)^2^ nm^2^, the sarcomere force 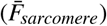 was calculated as:

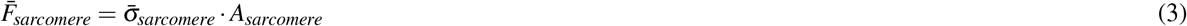

### Statistical Analysis

Combination of GraphPad Prism version 10 and R (R version 2024.4.2.764)^60^ were used to conduct statistical tests. Significance testing was done using Welch’s t-tests in Prism for univariate datasets (structural metrics, average sarcomeric force). A log-linked, generalized linear model with gamma distribution was applied to stress data, testing for difference and interaction between seeding density and cell sex followed by post hoc test with pairwise estimate marginal means (EMM). Statistically significance is assumed at p < 0.05.

## Acknowledgements

We thank the members of the Cardiovascular modeling laboratory at the University of California, Irvine, especially Nida T. Qayyum, for their useful suggestions throughout the preparation of this manuscript.

## Data availability statement

Raw data will be made available through Dryad repository. All data aggregates are within the manuscript.

## Author contributions statement

M.T. conducted the experiments and analyzed results. T.N. prepared experimental materials. S.K. contributed to result analyses for figures 1 and 2. M.T. and A.G. conceived the project, interpreted results, wrote and revised the manuscript. All authors reviewed the manuscript.

